# Neural representation of linguistic feature hierarchy reflects second-language proficiency

**DOI:** 10.1101/2020.06.15.142554

**Authors:** Giovanni M. Di Liberto, Jingping Nie, Jeremy Yeaton, Bahar Khalighinejad, Shihab A. Shamma, Nima Mesgarani

## Abstract

Acquiring a new language requires a simultaneous and gradual learning of multiple levels of linguistic attributes. Here, we investigated how this process changes the neural encoding of natural speech by assessing the encoding of the linguistic feature hierarchy in second-language listeners. Electroencephalography (EEG) signals were recorded during English story listening from native Mandarin speakers with varied English proficiency and from native English speakers. We measured the temporal response functions (TRF) for acoustic, phonemic, phonotactic, and semantic features in individual participants and found a main effect of proficiency on the linguistic encoding. This effect of second-language proficiency was particularly prominent on the neural encoding of phonemes, showing stronger encoding of “new” phonemic contrasts (i.e. English contrasts that do not exist in Mandarin) with increasing proficiency. Overall, we found that linguistic feature representation in nonnative listeners progressively converged to that of native listeners with proficiency, which enabled accurate decoding of language proficiency. This detailed view advances our understanding of the cortical processing of linguistic information in second-language learners and provides an objective measure of language proficiency.

## Introduction

Instructed second-language (L2) learning is a time-consuming and challenging process. Adult learners rarely attain native-like L2 proficiency and instead carry-over features of their native languages to their L2^1–3^, which can have a major impact on their social lives^4–6^. Listening to L2 speech elicits stronger neural activations in shared linguistic areas^7,8^ and engages cortical areas that are not active when listening to the native language^9^. However, there remains considerable uncertainty on the precise neural changes that underpin the increased L2 proficiency that come with the learning process^8,10,11^.

To elucidate the neural mechanisms that underlie L2 perception, it is crucial to assess the effect of proficiency on objective neural measures that capture the multifaceted cortical encoding of language. This is a complex task especially because speech perception involves the analysis of various acoustic and linguistic features, a process that is thought to engage a hierarchical neural network composed of various interconnected cortical regions^12^. Distinct stages of processing were shown to be affected differently by proficiency, with some of them becoming more native-like than others for proficient L2 users. Part of the evidence comes from electro- and magneto-encephalography (EEG and MEG respectively) research, which showed the effect of proficiency at the levels of phonemes^13^, syntax^14,15^, and semantics^16^. These studies measured the changes in well-known event-related potentials, such as the MMN, N400, and P600. These approaches, however, use unnatural speech stimuli (e.g., isolated syllables or violative speech sentences) which do not fully and realistically activate the specialized speech cortex^17–19^. In addition, these approaches consider various levels of speech perception independently and in isolation. Language learning, on the other hand, involves simultaneous acquisition of novel phonetic contrasts^20,21^, new syllabic structures (phonotactics)^22^, and new words. A more complete view of the neural basis of language learning therefore requires a joint study of multiple levels of the linguistic hierarchy to advance our understanding of L2 perception by informing us on the precise effect of proficiency on the cortical processing strategies that underpin sound and language perception^23–25^.

Previous effort in using naturalistic speech stimuli to study language proficiency showed a modulation of EEG phase-synchronization during naturalistic speech listening both at sub-cortical (FFR^26,27^) and cortical levels (gamma EEG synchrony^28,29^). Specifically, stronger synchrony between EEG channels was shown for low-proficiency users^29^, which is in line with theories positing that less experienced listeners require stronger cortical engagement, such as the *cortical efficiency theory*^29,30^. However, that work could not isolate neural signatures at particular linguistic stages. Recent studies have successfully isolated neural signatures of various linguistic levels based on speech-EEG synchrony (cortical tracking^31^) from a single electrophysiological recording. Such measures were derived based on low-frequency cortical responses to natural speech from audio-books^32–34^ and cartoons^35,36^, which were recorded non-invasively on both children and adults. Here, we adopted that framework to investigate how proficiency shapes the hierarchical cortical encoding in L2 subjects, and how that differs from L1 subjects. Our analysis focused on speech processing at the levels of sound acoustics^37,38^, phonemes^32,34^, phonotactics (statistics on phoneme sequences^33,39^), and semantics^40–42^. We hypothesized that the neural encoding of all three levels of linguistic properties would be modulated by L2 proficiency, becoming more native-like without fully converging^43,44^. A different progression of this learning effect was expected for distinct linguistic levels. Specifically, we predicted phoneme and phonotactic responses, which benefit from but do not require sentence comprehension, to show a continuous progression starting from the earliest stages of learning, partly as a form of implicit learning^45^. Semantic-level encoding was also expected to show progressively stronger encoding^46^, however with a most prominent change at an intermediate level of proficiency, when the comprehension of few words facilitates the understanding of the neighboring ones (e.g., semantic priming^47,48^), thus determining a turning point beyond which comprehension increases drastically.

To shed light on the neural mechanisms underlying the encoding of linguistic features, the present study combines objective neural indices of acoustic and linguistic processing to assess the differences between L2 subjects with varying proficiency levels during a natural speech listening task. We expected the hierarchical linguistic encoding in L2 participants to change with increased proficiency, becoming progressively more similar to the neural linguistic encoding of native speakers.

## Results

62-channel EEG was recorded from 51 participants as they listened to continuous speech sentences in English. Participants were native Standard Chinese (Mandarin) speakers with English as a non-native language (L2 listeners). Two of these subjects were excluded because of synchronization issues with the EEG recordings. English proficiency was assessed by means of a standardized test of receptive skills that assigned participants to six different CEFR levels (Common European Framework of Reference for languages): *basic user* (A1 and A2 levels), *independent user* (B1 and B2 levels), and *proficient user* (C1 and C2 levels). The recruitment of participants continued until 17 participants were identified for each A, B, and C group (**Supplementary Figure 1B**).

Subjects listened to 1.5 hours of speech material taken from a children’s story book (Hank the Cowdog) narrated by four speakers (2 female). The gender of the speaker alternated for each consecutive sentence. The experiment was divided into twenty blocks, each about three minutes long, grouped in five sections (each composed of four blocks). The experiment included behavioural tasks to monitor the engagement to the speech material for participants with different language proficiency levels (**Supplementary Fig. 1C,D**). The first test was a word comprehension task, where at the end of each experimental block subjects were asked to select which words had been spoken in the last block from a list of eight. The second task, which did not require comprehension, was a gender identification task, where the subjects were asked to report the gender of the speaker that uttered the last sentence before the end of the experimental block. Finally, subjects did a one-back repetition detection task in which they were required to detect repeated phrases (containing 2-4 words and was always at the end of a sentence), which occurred 1-5 times per experimental block. Participants were instructed to use a clicker counter when they heard a repeated phrase and to report the number of clicks at the end of each block. EEG data corresponding to the phrase repetition were excluded from the analysis to remove contamination due to the motor action. The detection of repeated sounds has been successfully used in many studies using non-speech and nonsense speech sounds ^49–51^. The design and result of the behavioral tasks is shown in **Figure 2A,B**. As expected, the word comprehension score increased significantly with proficiency (ANOVA, *F*(1.8, 44.3) = 24.1; *p* = 5.4*10^−8^, post-hoc comparisons: *p*_A-vs-B_ = 0.003, *p*_B-vs-C_ = 0.084, *p*_A-vs-C_ < 0.001). However, all subjects were able to perform the gender identification and one-back tasks with similar accuracy across proficiency levels, suggesting a comparable degree of engagement among participants across groups (*gender identification task*: ANOVA, *F*(1.4, 34.7) = 0.1, *p* = 0.90; *one-back task*: ANOVA, *F*(1.9, 45.9) = 1.1; *p* = 0.34).

**Figure 2.**
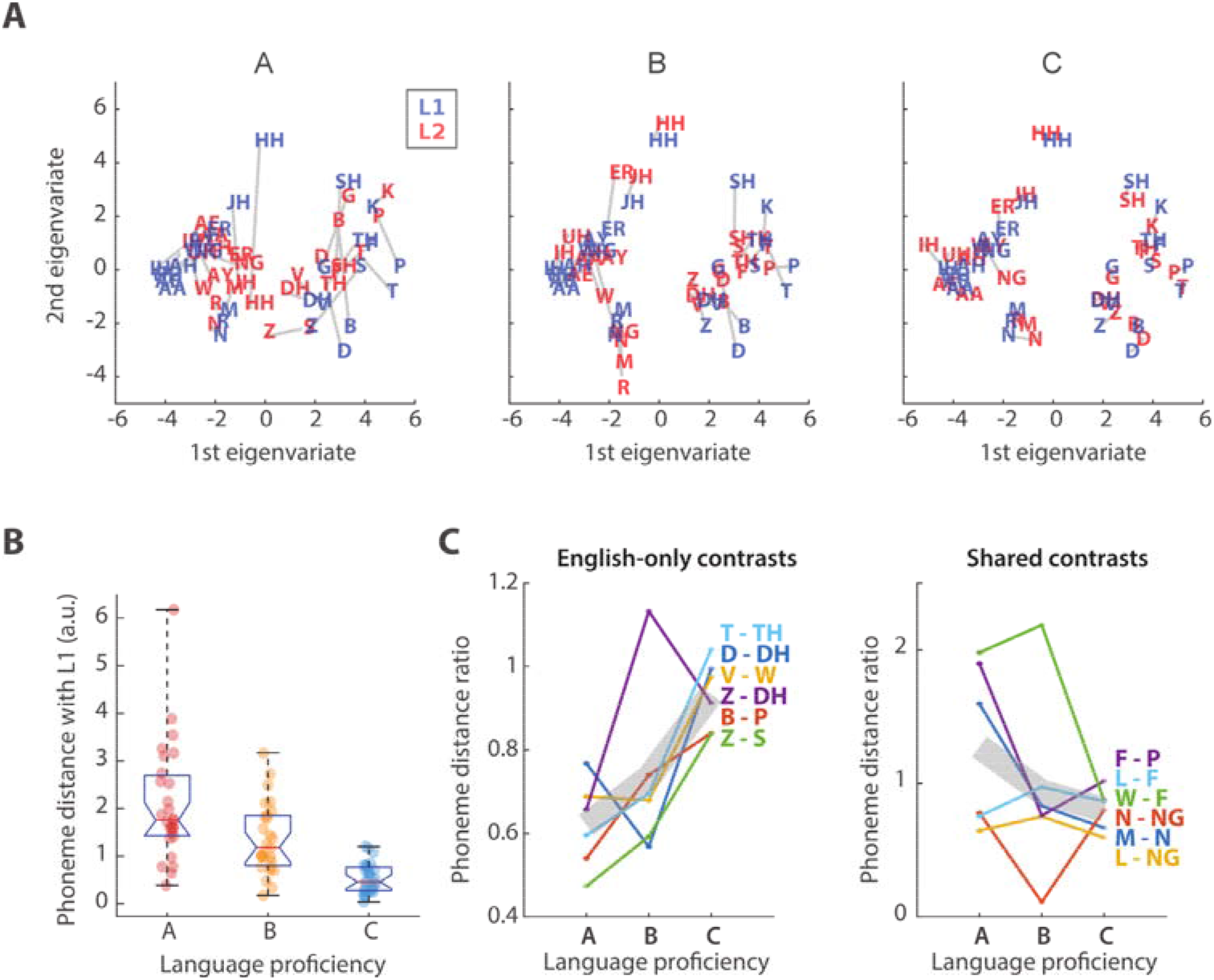
Effect of proficiency on L2 phoneme encoding. (A) Phoneme distances for L1 and L2 are compared for each proficiency group by means of a multi-dimensional scaling (MDS) analysis based on the TRF_Ph_ weights at the electrode Cz and peri-stimulus time-latencies from 0 to 600 ms. Blue and red colors indicate phonemes for L1 and L2 participants respectively. (B) Distance between L1 and L2 phonemes for each language proficiency group. A significant effect of proficiency was measured on the L1-L2 phoneme distance (ANOVA, p = 1.6*10^−8^). Error bars indicate the SE of the mean across phonemes. (C) Distance between phoneme-pairs for each proficiency level. The left panel shows results for contrasts existing in English but not in Standard Chinese, where we expected increasing discriminability with proficiency due to learning. The right panel shows distances for contrasts that exist both in English and Standard Chinese, where we did not expect a learning effect. Values were divided by the distance for L1 participants. Grey lines indicate the mean across all selected phonemic contrasts.

The analyses that follow aim to test the hypothesis that proficiency in a second language changes the neural tracking of linguistic properties, to demonstrate that non-invasive EEG is sensitive to those changes, and to determine its sensitivity to that change at the level of individual subjects and linguistic properties. We were particularly interested in testing if the neural encoding of individual phoneme contrasts could be assessed and, if so, whether proficiency improved the neural encoding of the “new” phoneme contrasts, i.e. that exist in English but not in Standard Chinese (the native language of the L2 listeners). We also aimed to relate the effect of L2 proficiency with previous work on natural speech perception. To do so, we analysed EEG data from a previous experiment^34^, which was collected from 22 native English speakers (L1 listeners) as they listened to the same continuous English speech stimuli presented to the L2 group in the present experiment. While this comparison is useful to put the results into the context of the previous literature, our analyses primarily tackle the effect of proficiency within the L2 group.

To investigate the low- *versus* higher-level brain processing of speech, we used linear regression models to measure the coupling between the low-frequency cortical signals (1–15 Hz) and progressively more abstract properties of the linguistic input. Specifically, we selected features describing acoustic properties based on the speech envelope (amplitude envelope – **Env**, spectrogram – **Sgr**, half-way rectified first derivative of the envelope **Env’**^52,53^), phoneme onsets (**Pon**), phonetic articulatory features (**Phn**^20,21^), its reduced version marking only vowels and consonants (**Pvc**), phonotactics (**Pt**^54^), and semantic dissimilarity (**Sem**^55,56^). Then, we used multiple linear regression to estimate the temporal response function (TRF) describing the spatio-temporal relationship between stimulus descriptors and EEG signal (**Figure 1**^57–59^). The combination of multiple speech features in a single multivariate model allowed us, for the first time, to assess the hierarchical processing of L2 speech from a single EEG recording session based on natural speech.

**Figure 1.**
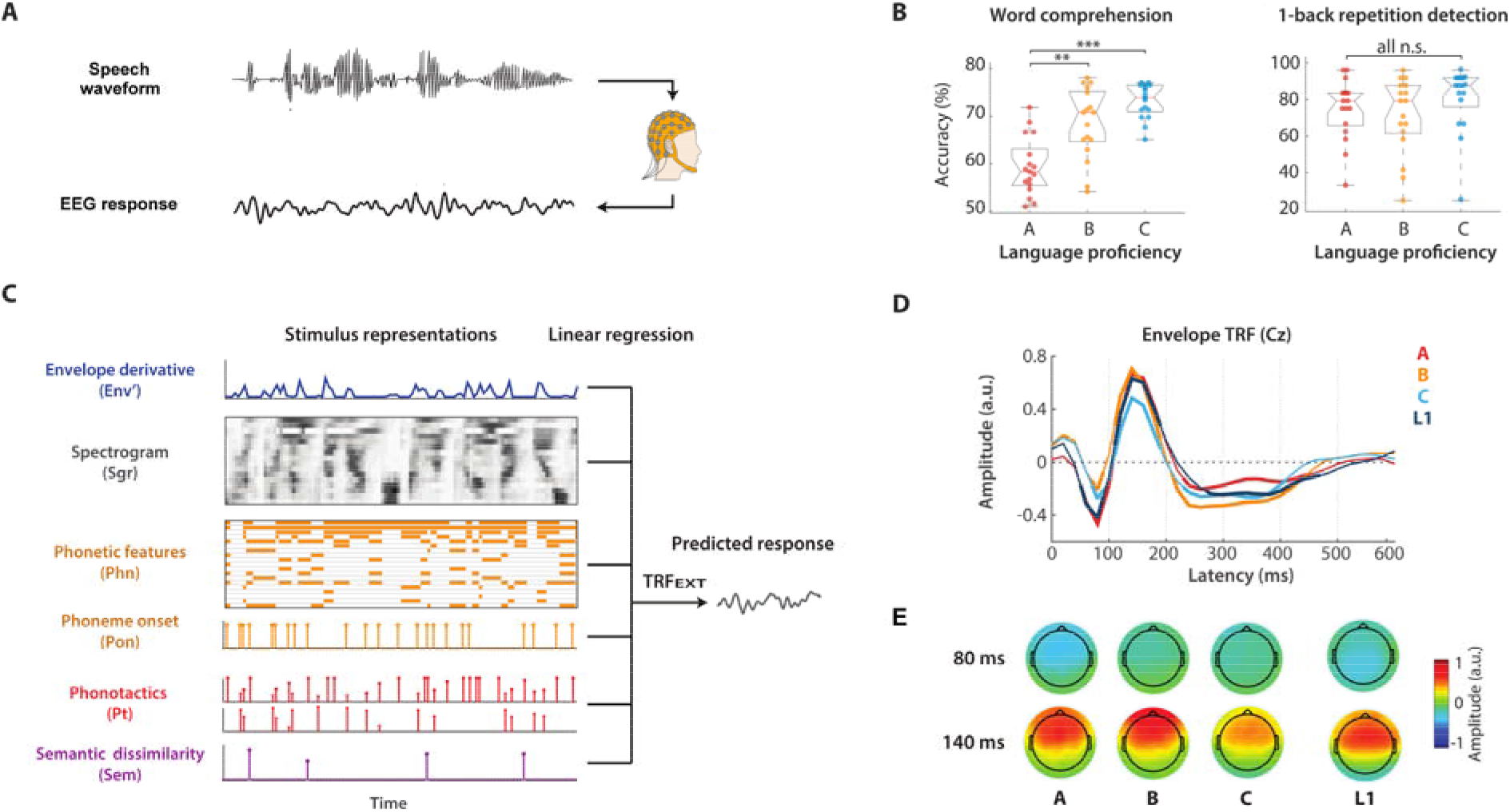
Investigating the hierarchical cortical encoding of language with the Temporal Response Function (TRF) analysis framework. (A) Multichannel EEG signal was recorded as participants listened to audio-stories. Participants were asked to press a clicker counter when detecting a one-back phrase repetition (2-4 words), which occurred 1-5 times per experimental block. At the end of each block, participants were asked to report the number of repetitions, to identify words that were spoken during the block from a list of eight, and to indicate the gender of the speaker. (B) Results for the word comprehension and the 1-back detection task. Significant group differences (ANOVA, ** p<0.01, *** p < 0.001) were measured for the ‘word comprehension’ score that positively correlated with proficiency. No significant effects emerged for ‘one-back tasks’ and ‘gender identification’ as they are independent from the proficiency levels. The gender identification result is not reported as accuracy was larger than 95% for all participants. (C) Acoustic and linguistic information were extracted from the stimulus and encapsulated into data vectors and matrices. Multivariate linear regression was used to identify a linear fit that optimally predicted the EEG signal from features at multiple linguistic levels (EXT). The same procedure was also run on a more compact set of descriptors (ALL), which differed in that Sgr and Phn were replaced by Env (broadband envelope instead of a 16-band spectrogram) and Pvc (indicator variables for vowel and consonants only rather than for a 19-dimensional set of phonetic features) respectively. (D) Envelope TRF weights (TRF_Env_) averaged across all EEG channels at peri-stimulus time-latencies from 0 to 600 ms. TRF_Env_ was part of a model that was fit by including features at all other levels of interest (ALL). Thick lines indicate weights that were statistically different from zero across all subjects of a group (p < 0.05, FDR corrected permutation test). (E) Topographies of the TRF weights across channels for two selected time-latencies.

### Hierarchical cortical encoding of non-native speech

Forward TRF models were fit between concatenations of speech features and the EEG signal for a broad time-latency window from 0 to 600ms to take into account various time dependencies at all hierarchical levels of interest. The stimulus descriptor included **Env**, **Env’**, **Pon**, **Pvc**, **Pt**, and **Sem**(**ALL;** see **Methods**). This combination of features allowed us to capture and discern EEG variance corresponding to various hierarchical stages, while using a low-dimensional descriptor (8 dimensions). We also fit TRF models with an extended stimulus descriptor (**EXT**) that includes **Sgr**, **Env**’, **Pon**, **Phn**, **Pt**, and **Sem**, which provided us with a higher level of detail in the spectrotemporal and phonological processing of speech. However, this increased dimensionality of the model (40 dimensions) makes fitting the model more challenging. Leave-one-out cross-validation indicated that the resulting TRF models could reliably predict the EEG signal for all subjects (*r_ALL_* > r_ALL_SHUFFLE_ and *r_EXT_* > r_EXT_SHUFFLE_, *p* < 0.01, permutation test where input sentences were randomly shuffled, N = 100; EEG prediction correlations were averaged across all electrodes).

**Figure 1D,E** show the model weights corresponding to the **Env** descriptor (part of TRF_ALL_) after averaging across all electrodes and all subjects within each proficiency group (A, B, C, and L1). TRFs for the four groups appear temporally synchronized, which was expected for cortical responses to low-level acoustics. Effects of proficiency emerged on the TRF_Env_ magnitude that, however, did not survive correction for multiple comparison at any of the EEG channels (two-tailed permutation tests with FDR-correction over time-latencies, N = 100). In spite of that, the envelope response in L2 participants could have plausibly become more similar to that of native speakers with proficiency. To test for this possibility, we measured the Pearson’s correlation scores between the TRF_Env_ for each L2 subject and the average TRF_Env_ weights across all L1 participants. This measure of similarity between L1 and L2 did not show any significant difference between A, B, and C groups (*p* > 0.05).

Although envelope TRFs have been proven robust and contributed to the study of various aspects of auditory perception^60–63^, we have also modeled the low-level auditory responses by considering the acoustic spectrogram (**Sgr**), which was shown to be a better predictor of the EEG signal^32,64^. However, observing TRF_Sgr_ (part of TRF_EXT_) for different auditory frequency bands did not lead to new clear-cut insights in this case, thus the rest of the manuscript will focus on the envelope TRF results.

#### Effect of proficiency on the cortical encoding of phonemes in L2 listeners

Phonetic feature information was represented by the categorical descriptor **Phn**, which marked the occurrence of a phoneme with a rectangular pulse for each corresponding phonetic features (see **Methods**)^32^. TRFs were fit for each subject by combining the **Phn** descriptor with all others in the **EXT** feature-set. The weights corresponding to the descriptor of interest, TRF_Phn_, were extracted from TRF_EXT_. In this case, the other descriptors played the role of nuisance regressors, meaning that they reduced the impact of acoustic-, phonotactic- and semantic-level responses on TRF_Phn_. The effect of proficiency was assessed in L2 participants by measuring the change in TRF_Phn_ between the proficiency levels A, B, and C. First, TRF_Phn_ was rotated from feature to phoneme space by means of a linear transformation (see **Methods**; **Supplementary Figure 2**). Then, a classical multidimensional scaling (MDS) was used to project the TRF_Phn_ (phonemes were considered as objects and time-latencies as dimensions) onto a 2D component space for each proficiency group (**Figures 2A**) where distances represent the discriminability of particular phonetic contrasts in the EEG signal. The result for each L2 proficiency group was then mapped to the average L1-MDS space by means of a Procrustes analysis. This analysis allows us to project the L2 phoneme maps for different proficiency levels to a common multidimensional space where they can be compared (the effect can be also measured, for example, by projecting A and B data to the C-level phoneme map). **Figure 2A** shows the average L1-L2 distance across all phonemes for each L2 participant, with blue and red fonts indicating phonemes for L1 and L2 participants respectively. Shorter L1-L2 distances were measured for increasing L2 proficiency levels (**Figure 2B**: ANOVA, *F*(1.4, 54.1) = 22.8; *p* = 1.6*10^−8^), indicating an effect of proficiency on the TRF_Phn_, with a progressive convergence of the phoneme TRF maps to that for native listeners.

To test whether the effect of proficiency on the phoneme maps was more pronounced for English phonemes that do not exist in Standard Chinese (the native language of L2 subjects), we measured the discriminability between selected pairs of phonemes based on TRF_Phn_ for each proficiency group. The phonemic discriminability was assessed by calculating the L2-L1 distance of a given phonemic contrast for each proficiency group A, B, and C. Distance values for each phoneme pair were normalized based on the L1 map for visualization. Using phonemes that can occur in minimal pairs (words differentiated by only one phoneme, e.g., “bat” /bæt/, “pat” /pæt/), and thus must be discriminable for proper comprehension, we selected ‘English-only contrasts’, which contain at least one phoneme that does not occur in Standard Chinese (e.g., TH). Such unknown phonemes have been shown to be perceived by L2 speakers as the closest existing phonemic neighbor in their L1, thus presenting challenges in discrimination^65,66^. We expected discriminability to increase with proficiency when considering phonemic contrasts that exist in English but not in Standard Chinese, thus reflecting the improved discrimination skills of L2 listeners. Our data is sensitive to this learning process, as we show in **Figures 2C** for six selected English-only contrasts (T vs. TH, D vs. DH, V vs. W, Z vs. DH, B vs. P, Z vs. S) which all show increased discriminability when comparing the A and C proficiency-level groups. As expected, this was not the case for phonemic contrasts belonging to both English and Standard Chinese languages (F vs. P, L vs. F, W vs. F, N vs. NG, M vs. N, L vs. NG) which did not show any consistent change with proficiency.

#### Proficiency modulates phonotactic responses at both short and long latencies

In a given language, certain phoneme sequences are more likely to be valid speech tokens than others. The likelihood of a phoneme sequence *p_1..n_* being a valid speech token can be estimated with statistical models based on language-specific rules. One such model, called BLICK^54^, returns the phonotactic *score* corresponding to the negative logarithm of the sequence likelihood. As such, that score has larger values for less likely (more surprising) sequences, providing us with a quantitative comparison between speech tokens. Crucially, this score is sensitive to changes in likelihood between the valid words composing natural conversational speech, thus allowing us to go beyond previous research based solely on phonotactic violations. Here, likelihood values for each phoneme at position *k* of a word were calculated as the phonotactic score on each subsequence *p*_1..k_, and used to modulate the amplitude of a phoneme onsets vector (all other time points were assigned to zero^39^; see **Methods**). **Figure 3A** compares the corresponding TRF weights (part of TRF_ALL_) between proficiency groups at three scalp locations of interest. Qualitatively different TRF patterns emerged between groups, with an early positive component (~40 ms) that emerged consistently for all groups, an expected longer latency component (300-500 ms) that was less pronounced for L2 than L1 subjects, but it was significant for L2 with high and medium proficiency, and an unexpected earlier component (~120 ms) that emerged consistently only for all L2 groups but not L1 (FDR-corrected permutation tests). The same latencies showed significant effects of proficiency, which was measured as a Pearson’s correlation between proficiency level and TRF magnitude. The topographical patterns in **Figure 3B** further clarify that this effect of proficiency was distributed across most scalp areas, but especially in centro-frontal scalp areas at 120 ms, while the effect at a latency of about 360 ms showed centro-parietal patterns. We also assessed the distance between the TRF of each L2 participant and the average TRF across all L1 participants. This distance, which was calculated with a cosine metric over all electrodes and time-latencies of the TRF, did not show a significant effect of proficiency (**Figure 3C**: ANOVA, *F*(1.4, 33.2) = 2.0; *p* = 0.14), indicating that the effect of proficiency measured at individual TRF components were not sufficiently pronounced to emerge when considering all channels and latencies.

**Figure 3.**
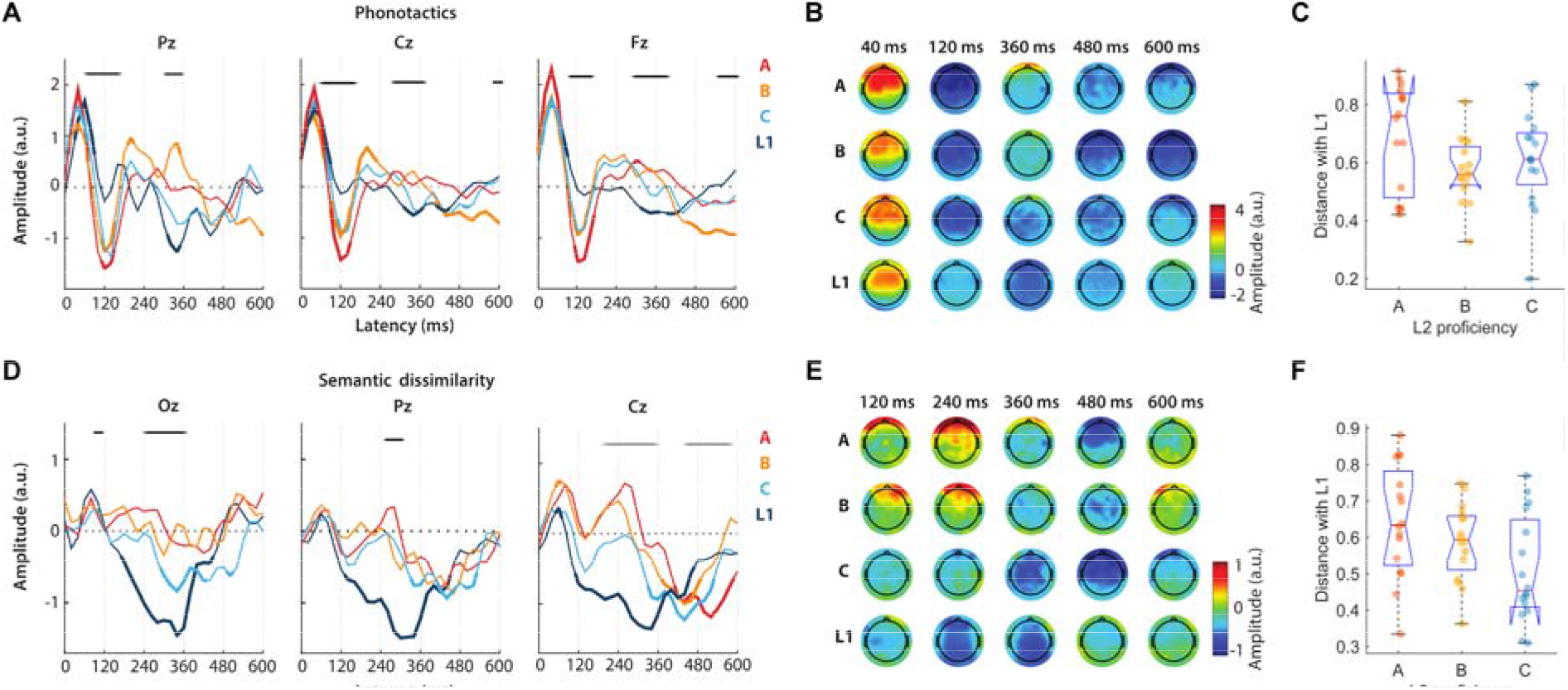
Effect of proficiency on the EEG responses to phonotactics and semantic dissimilarity regressors. (A) Model weights of the phonotactic TRF for three selected midline EEG channels at peri-stimulus time-latencies from 0 to 600 ms. Results for distinct participant-groups are color-coded. Thick lines indicate weights that were statistically different from zero across all subjects of a group (p < 0.05, FDR corrected permutation test). Horizontal black lines indicate significant correlations between TRF weights and language proficiency (p < 0.05, Pearson’s correlation). (B) Topographies of the phonotactics TRF weights for five selected time-latencies. (C) Cosine distance of the phonotactics TRF for individual L2 participants with the average L1 TRF. The distance was calculated based on all electrodes and time-latencies. (D) Model weights of the semantic dissimilarity TRF for selected EEG channels. (E) Topographies of the semantic dissimilarity TRF weights for five selected time-latencies. (F) Cosine distance of the phonotactics TRF for individual L2 participants with the average L1 TRF.

#### Stronger and earlier cortical responses to semantic dissimilarity with proficiency

A similar analysis was conducted based on semantic dissimilarity rather than phonotactic scores. Specifically, content words were described according to a 400-dimensional feature space that was identified based on word co-occurrence (Word2Vec algorithm^55^). Then, semantic dissimilarity was quantified as the *distance* of a word with the preceding semantic context, thus resulting in a vector marking the onset of all content words with these distance values (see **Methods**)^40^. **Figure 3D** shows the semantic dissimilarity TRF_Sem_ (part of TRF_ALL_) for three selected scalp channels. Average TRF_Sem_ for L1 participants were consistent with the results shown by Broderick and colleagues^40^, with a negative component peaking at peri-stimulus latencies of 340-380 ms. Similar TRF patterns emerged for the L2 C-level participants, whose average TRF_Sem_ showed a negative component at comparable time-latencies, with trough at latencies between 340 and 440 ms (depending on the EEG channel). As expected, we observed significant correlations between TRF magnitude and proficiency over central and posterior scalp areas (**Figure 3E;***p* < 0.05, Pearson’s correlation). Interestingly, an unexpected significant bilateral centro-frontal negativity (BCN) peaking between 440 and 520 ms appeared in all L2 subjects but not in L1 subjects. As for phonotactics, we also assessed the distance between the TRF of each L2 participant and the average TRF across all L1 participants. In this case, this distance showed a significant effect of proficiency (**Figure 3F**: ANOVA, *F*(1.83, 42.1) = 3.7; *p* = 0.033), indicating a robust progressive L2-to-L1 convergence of the semantic dissimilarity TRFs with proficiency.

### Decoding language proficiency

Our results indicate that language proficiency modulates the cortical responses at various linguistic processing levels. Given this relation, we examined the extent to which the proficiency of a subject could be predicted from the combined effects of different linguistic features. First, multilinear principal component analyses (MPCA) was conducted on the TRF weights corresponding to **Env**, **Phn**, **Pt**, and **Sem** separately, and the first component was retained for each of them. In doing so, information spacing along three dimensions (EEG channels, time latencies, and stimulus features – e.g., phonetic features) was compressed into a single value for each participant. A linear regression model was then fit to predict L2 proficiency (L1 subjects were excluded from this analysis) based on those four TRF features. **Figure 4A** shows the effect of each regressor on the model fit (coefficient estimate and standard error), with an overall regression correlation *r* = 0.68. Significant effects were measured for each of the four features, and this was true also when the ‘age’ information and the ‘one-back repetition detection’ score (which was a measure of the attentional engagement to the experiment) were included in the regression fit. This result confirmed that the main effect of proficiency was not due to attention nor age.

**Figure 4.**
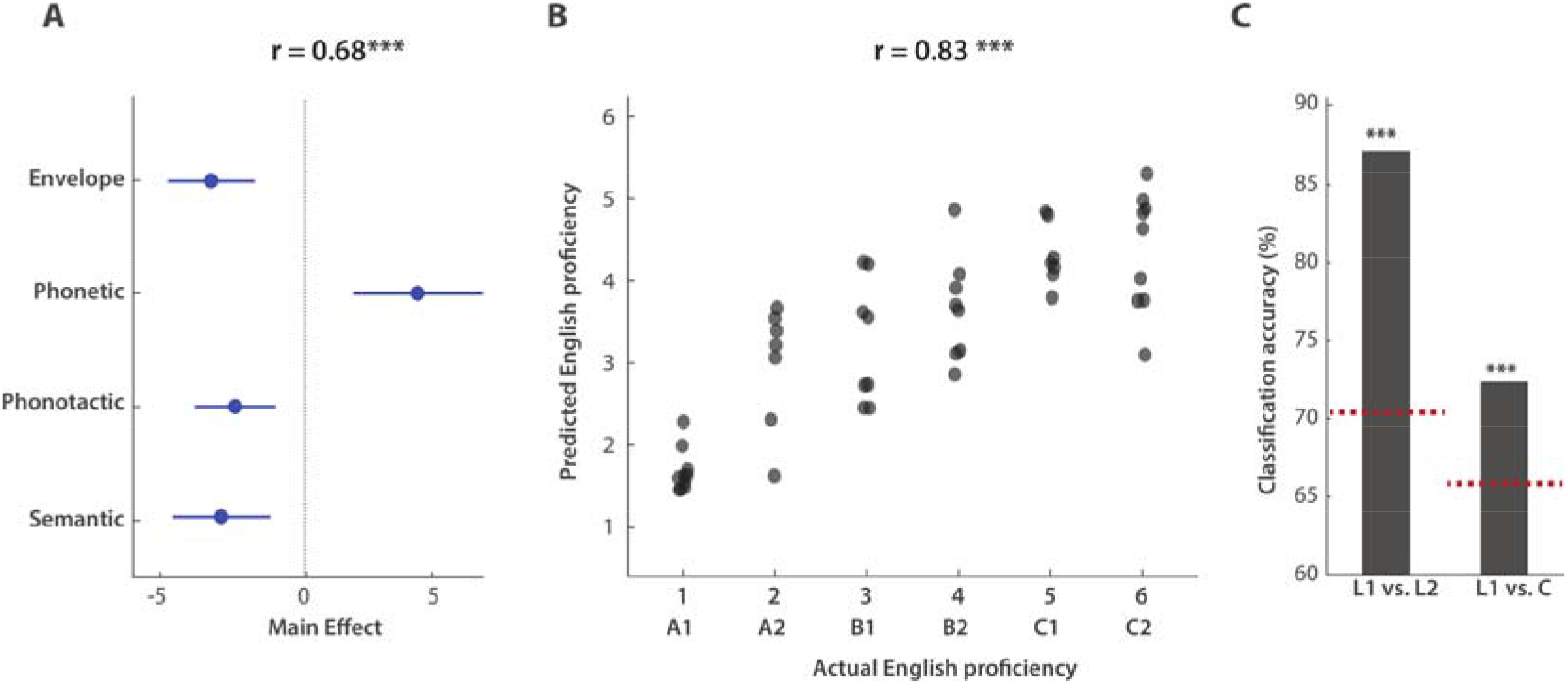
Accurate decoding of L2 proficiency from EEG data. (A) A multilinear principal component analysis (MPCA) was performed on the TRF weights corresponding to speech descriptors at all linguistic levels of interest. The first MPCA component was retained for the TRFs corresponding to Env, Phn, Pt, and Sem. The combination of those four features was predictive of L2 proficiency (r = 0.68), with significant effects for all features which were not due to group-differences in age or attention. (B) A support-vector regression analysis shows that EEG data accurately predict the L2 proficiency level at the individual subject level (r =0.83, MSE = 1.14). (C) Classification accuracy for L1 versus L2 and L1 versus C-level L2. The red dotted lines indicate the baseline classification levels, which were calculated as the 95^th^ percentile of a distribution of classification accuracies derived after random shuffling the output class labels (N = 100).

A similar decoding approach was then used to assess whether and how robustly L2 proficiency could be decoded based on EEG indices of language processing. A set of 26 features was identified to most comprehensively describe the effects of L2 proficiency on the TRFs. Features were based either on the TRF weights (as for **Figure 4A**), on the EEG prediction correlations based on subject-specific TRF models, or on EEG prediction correlations for each subject when using average TRF models that were fit on the other subjects, for A, B, C, and L1 groups separately (generic modelling approach^36,67^) (see **Methods** for a detailed list of features). Each of the 26 feature-vectors had multiple dimensions (e.g. electrodes, time-latency). For this reason, as it was described above, MPCA was used to reduce those vector to one-dimensional regressors. Support Vector Machine (SVM) regression was used to decode L2 proficiency based on such regressors. A backward elimination procedure identified a reduced set of features (**Supplementary Table 1**) whose combination produced optimal L2 proficiency decoding scores, with *MSE* = 1.14 and Pearson’s correlation *r* = 0.83, *p* = 3*10^−13^ (**Figure 4B**). Another way to quantify the quality of the proficiency decoding is to assess the A- vs. C-level classification by placing a simple threshold on the prediction values (value 3.5, which cuts the prediction space in half). This binary classification could identify A- vs. C-level participants with 91% accuracy.

Further analyses were conducted to assess the effect of “nativeness” on the EEG responses to speech. Specifically, differences in language processing between L1 and L2 subjects may be in part driven by a fundamental distinction between native and non-native language processing that is not due to proficiency *per se,* but rather due to differences in the L1 and L2 processing networks^68,69^. In fact, the TRF results in **Figures 1–3** indicated that higher proficiency levels do not always lead to EEG responses that are equivalent to the ones of native speakers. Specifically, while there was some level of L1-L2 convergence for phoneme-level TRFs, this phenomenon was less pronounced for phonotactics and semantic dissimilarity responses, with marked differences between L1 and C-level L2 (e.g., the latency of the negative component at ~120 ms in TRF_Pt_). Here, we attempted to disentangle those differences from the effect of L2 proficiency by conducting an SVM binary classification analysis for L1 versus L2 participants. This procedure used the same 26 features and backward elimination strategy as in the previous regression analysis. First, an L1 versus L2 classification accuracy of 87% was obtained when all 71 subjects were included in the analysis, with a baseline classification accuracy of 70% (95^th^ percentile of a distribution of classification accuracy values when L1-L2 labels were randomly shuffled – 100 shuffles). In order to reduce the contribution of proficiency to the classification result, the same analysis was run on L1 and C-level L2 participants only. In this case, a classification accuracy of 73% was measured, with a baseline of 66%, thus suggesting that the EEG responses to continuous speech reflect both the influence of L2 proficiency and nativeness. Nevertheless, it is important to highlight that this result emerged on a small cohort of L1 and L2 participants. Furthermore, different behavioural tasks were used for L1 and L2 participants as the L2 group included subjects that could not understand the speech. As such, further work with a more specific design and a larger sample size is needed to confirm this result.

## Discussion

The human brain responds differently when listening to second-language and native speech^13,15,16^, typically leading to lower listening performances that vary between individuals and can be quantified with standardized language tests. Despite the general consensus for the cognitive, social, and economic advantages that come with high L2 proficiency, the neural underpinnings of second language perception and learning remain unclear^70,71^. One reason why this issue remains unresolved is methodological. Experimental evidence derived from direct neural measures is minimal and often limited to single linguistic properties^72–76^, thus offering only a partial view of this complex brain mechanism. The present study establishes a methodological framework to provide a more comprehensive examination of the language processing system in naturalistic conditions. We isolated neural indices of speech perception at multiple processing stages from EEG responses to natural speech, revealing marked effects of L2 proficiency that were robust at the individual subject level. Overall, the results confirm our hypothesis that the cortical encoding of speech in L2 listeners changes with proficiency and that EEG responses to natural speech are sensitive to its change for distinct linguistic properties and even at the level of individual phonemic contrasts.

Previous studies that investigated L2 perception in naturalistic paradigms focused on the relationship between neural activity and acoustic envelope, and found stronger coupling in L2 than L1 subjects^77^. That EEG result, which was found with a selective attention listening task in a multi-talker scenario, pointed to a link between increased listening effort and stronger cortical tracking of the speech envelope. However, it remained unclear which of the linguistic and non-linguistic properties of speech correlate with the acoustic envelope results in this increased cortical tracking. In fact, an increased coupling between EEG and speech envelope could reflect increased encoding of acoustic features, stronger reliance on higher order processes, or even activation of distinct cortical areas. For example, recent work indicated that envelope tracking increases with age due to a stronger engagement of higher order areas, thus reflecting a difference in processing strategy for older listeners^78,79^. While our results did not show differences in envelope tracking (EEG prediction correlations) with proficiency, the shape of the envelope TRF significantly contributed to L2 proficiency decoding (**Figure 4**), which is in line with a link between acoustic-level encoding and effort. Interestingly, this result was obtained on a single-talker task with no competing noise. Therefore, the application of this same approach on a more cognitively demanding task^62,80^ could help tease apart the effects of L2 proficiency and listening effort on the cortical encoding of acoustic features.

As we had hypothesized, the cortical encoding of phonemes changed with L2 proficiency, becoming progressively more similar to L1, which is in line with perceptual theories such as the expanded Native Language Magnet Theory (NML-e^81^) and the Perceptual Assimilation Model (PAM-L2^82^). Our TRF analysis has discerned individual phonemic contrasts, showing that the cortical encoding of phonemes becomes progressively more sensitive to contrasts existing in English but not Standard Chinese (**Figure 2**). This work extends previous findings on the cortical encoding of phonemes^32,34,64^ by demonstrating that EEG responses to natural speech show sensitivity to individual phoneme contrasts, with response patterns that become progressively more categorical with proficiency. Furthermore, that result goes beyond previous work^64^ by revealing a low-frequency EEG component that could not be explained by simple acoustic features such as acoustic envelope, derivative of the envelope, and spectrogram. Our results are in line with the majority of theories on L2 perception which suggest the impact of a subject’s L1 on phonological encoding of the L2. Specifically, **Figure 2A** indicates that the native language constitutes a “starting point” for the phonological encoding in L2 learners, which then changes with experience and converges toward the encoding for L1 listeners. Reproducing this work on participants with other native languages could provide us with detailed insights on the effect of the native language on the phoneme encoding in high proficiency L2 learners. Furthermore, additional data based on a more balanced design, with subjects listening to both their native and non-native language, could reveal whether and how learning a particular L2 influences the cortical processing of the native language^19,83^, as was postulated by the bidirectional cross-linguistic influence principle in the Speech Learning Model (SLM^84^).

Proficiency was also shown to shape language encoding at the level of phonotactics, with TRFs in L2 subjects progressively converging toward L1 TRFs. Our results indicate two effects of phonotactics. First, an hypothesized, TRF component peaking at speech-EEG latencies of about 300-450ms, as measured in a previous EEG study by our group^39^, with more negative responses for higher proficiency-levels (**Figure 3A**); Second, an effect at shorter latencies of about 120ms, where a negative component that was not present for L1 participants emerged for L2 participants. Interestingly, a component reflecting phonotactics was previously measured at that speech-neural signal latencies with MEG^33^ but not EEG. Our finding provides a new link between the EEG and MEG literature by clarifying that phonotactics modulates EEG responses at both shorter and longer latencies, and that the effect at shorter latencies emerges for L2 learners but not L1. This discrepancy may be due to the difference in the type of signal recorded by EEG and MEG modalities. The larger values for lower-proficiency users could reflect an effect of surprise on the phoneme sequences due to the use of an incorrect (or imperfect) model of phonotactics.

Semantic dissimilarity TRFs were previously shown to be characterized by a negative centro-parietal component at speech-EEG latencies of about 350-400ms. This finding is in line with previous work on the N400 event-related response^42,85,86^ which shows that this component is modulated by intelligibility and attention^40^. Similarly, we expected a strong response negativity for users with higher language proficiency, and no response for people with no English at all. Consistent with this hypothesis, our results identified a posterior component with magnitude that increases with proficiency (**Figure 3D**). In addition, an unexpected centro-frontal component arose at latencies of about 440-520ms which was negatively correlated with the latency of response rather than the magnitude of the component. This bilateral centro-frontal negativity (BCN) emerged even for participants with no English understanding, thus reflecting neural correlates time-locked to word onset but not semantics *per se*. This component may instead be related to other processes, such as the processing of sentence structure, memory, and learning of frequent words^87–90^. Further work is needed to clarify whether that signal reflects, for example, the familiarity with particular words, or it is related to ERP components such as the left anterior negativity (LAN), which was shown to reflect processing difficulties in morpho-syntax^91,92^.

Although both phonotactics and semantic level TRFs for L2 showed some level of convergence to L1, there was also a pronounced difference between L1 and C-level L2 participants, which was also reflected in the significant L1 versus C classification result in **Figure 4C.** This effect may reflect fundamental differences in the cortical mechanisms underlying L1 and L2 processing, rather than an effect of proficiency *per se*. This effect of nativeness that is somewhat different from the effect of proficiency is in line with the observation that a second language learnt after a certain critical (or sensitive) period usually leads to lower language proficiency than those of a native speaker^2,93,94^. More data could provide further insights on this topic, for example, by comparing L1 monolinguals with bilinguals and multilinguals with a wide range of learning-onsets of the English language. Further research is also needed to better understand the effect of nativeness, e.g. by comparing L1 and high-proficiency L2 listeners with a semantic task guaranteeing the same level of comprehension for all participants. Such a task could not be employed in the present study, whose primary focus was the effect of proficiency across L2 participants from A-to C-levels which, by design, presented variable levels of comprehension.

Our analysis focused on just few components of the speech processing hierarchy, namely the acoustic, phonemic, phonotactic, and semantic levels. One powerful element of this framework is that it can be extended to other levels of processing without the need for additional data. In fact, the EEG responses to natural speech likely reflect many more components of interest than the ones targeted in this occasion, and demonstrating how to isolate them would give us insights on each newly added feature and its link with proficiency, as well as giving us the chance to improve the accuracy of our EEG-based L2 proficiency assessment. For these reasons, we believe that a wide collaborative effort under a common protocol of data acquisition with EEG/MEG and natural stimuli could significantly and quickly advance our understanding of the speech and language cortical processing network (and could indeed extend to other questions of interest).

Understanding the neural underpinnings of second language perception and learning becomes particularly relevant when we consider that there are more children throughout the world that have been educated via a second (or a later acquired) language rather than exclusively via their L1^95^. Furthermore, there is evidence for perceptual advantage of bilinguals and multilinguals that is due to cross-language transfer^83,96,97^, and particular combination of languages may be better than others in the emergence of such a benefit. Further work in this direction may provide us with tools to predict the perceptual advantage that a particular second language would bring to a person given their background, thus constituting the basis for a procedure that, for example, could inform us on what second languages should be encouraged in school to particular individuals.

## Materials and Methods

### Participants

Fifty-one healthy subjects (twenty-four male, aged between 18 and 60, with median = 24 and mean = 27.5, forty-eight were right-handed) that learnt English as a second language (or that did not speak English) participated in the EEG experiment (L2 group). Two of these subjects were excluded because of issues with the EEG recordings (data could not be synchronized because of missing trigger signals). L2 participants were asked to take a 20-minute American English listening level test before the experiment. According to the results of this assessment, each participant was assigned to one of seven proficiency groups according to the CEFR framework (Common European Framework of Reference for Languages): A1, A2, B1, B2, C1, C2 (from low to high proficiency). A-, B-, and C- levels indicated basic, independent, and proficient users respectively. The A1 group included participants with no English understanding.

Data from twenty-two native English speakers (twelve male, aged between 18 and 45, twenty were right-handed) was originally collected for a previous study^34^ based on the same experiment, with the same setup and location (L1 group). All subjects reported having normal hearing and no history of neurological disorders. All subjects provided written informed consent and were paid for their participation. The Institutional Review Board of Columbia University at Morningside Campus approved all procedures.

### Stimuli and experimental procedure

EEG data were collected in a sound-proof, electrically shielded booth in dim light conditions. Participants listened to short stories narrated by two speakers (1 male) while minimizing motor movements and maintaining visual fixation on a crosshair at the center of the screen. The male and the female narrators were alternated to minimize speaker-specific electrical effects. Stimuli were presented at a sampling rate of 44,100 Hz, monophonically, and at a comfortable volume from loudspeakers in front of the participant. Each session consisted of 20 experimental blocks (3 min each), divided in five sections that were interleaved by short breaks. Participants were asked to attend to speech material from seven audio-stories that were presented in a random order. The engagement to the speech material was assessed by means of behavioural tasks. L2 participants were asked three questions at the end of each block. First, we asked whether the last sentence of the section was spoken by a male or female speaker. Next, participants were asked to identify 3-5 words with high-frequency in the sentence from a list of eight words. Third, participants performed a phrase-repetition detection task. Specifically, the last two to four words were repeated immediately after the end of some of the sentences (1-5 per block). Given that our target was monitoring attention, a finger-tip clicker was used to count the repetitions so that they would be engaged in a detection and not counting task, which would instead require additional memory resources and, potentially, reduce their engagement to the main listening task. Participants were asked to indicate how many sentences in the story presented these repetitions at the end of each block. To assess attention in L1 participants, three questions about the content of the story were asked after each block. All L1 participants were attentive and could all answer correctly at least 60% of the questions.

### EEG recordings and preprocessing

EEG recordings were performed using a g.HIamp biosignal amplifier (Guger Technologies) with 62 active electrodes mounted on an elastic cap (10 –20 enhanced montage). EEG signals were recorded at a sampling rate of 2 kHz. An external frontal electrode (AFz) was used as ground and the average of two earlobe electrodes were used as reference. EEG data were filtered online using a high-pass Butterworth filter with a 0.01 Hz cut-off frequency to remove DC drift. Channel impedances were kept below 20 kΩ throughout the recording.

Neural data were analyzed offline using MATLAB software (The Mathworks Inc). EEG signals were digitally filtered between 1 and 15 Hz using a Butterworth zero-phase filter (order 2+2 and implemented with the function *filtfilt*), and down-sampled to 50 Hz. EEG channels with a variance exceeding three times that of the surrounding ones were replaced by an estimate calculated using spherical spline interpolation.

### Speech features

In the present study, we have assessed the coupling between the EEG data and various properties of the speech stimuli. These properties were extracted from the stimulus data based on previous research. First, we defined a set of descriptors summarizing *low-level acoustic properties* of the music stimuli. Specifically, a time-frequency representation of the speech sounds was calculated using a model of the peripheral auditory system^98^ consisting of three stages: (1) a cochlear filter-bank with 128 asymmetric filters equally spaced on a logarithmic axis, (2) a hair cell stage consisting of a low-pass filter and a nonlinear compression function, and (3) a lateral inhibitory network consisting of a first-order derivative along the spectral axis. Finally, the envelope was estimated for each frequency band, resulting in a two dimensional representation simulating the pattern of activity on the auditory nerve^99^ (the relevant Matlab code is available at *https://isr.umd.edu/Labs/NSL/Software.htm*). This *acoustic spectrogram* (**S**) was then resampled to 16 bands^32,100^. A *broadband envelope* descriptor (**E**) was also obtained by averaging all envelopes across the frequency dimension. Finally, the *half-way rectified first derivative of the broadband envelope* (**E’**) was used as an additional descriptor, which was shown to contribute to the speech-EEG mapping and was used here to regress out the most acoustic-related responses as possible^64^.

Additional speech descriptors were defined to capture neural signatures of higher-order speech processing. The speech material was segmented into time-aligned sequences of phonemes using the Penn Phonetics Lab Forced Aligner Toolkit^101^, and the phoneme alignments were then manually corrected using Praat software^102^. *Phoneme onset* times were then encoded in an appropriate univariate descriptor (**Pon**), where ones indicate an onset and all other time samples were marked with zeros. An additional descriptor was also defined to distinguish between *vowels and consonants* (**Pvc**). Specifically, this regressor consisted of two vectors, similar to **Pon**, but marking either vowels or consonants only. While this information was shown to be particularly relevant when describing the cortical responses to speech^33^, there remains additional information on phoneme categories that contributes to those signals^32,103^. This information was encoded in a 19-dimensional descriptor indicating the *phonetic articulatory features* corresponding to each phoneme (**Phn**). Features indicated whether a phoneme was voiced, unvoiced, sonorant, syllabic, consonantal, approximant, plosive, strident, labial, coronal, anterior, dorsal, nasal, fricative, obstruent, front (vowel), back, high, low. The **Phn** descriptor encoded this categorical information as step functions, with steps corresponding to the starting and ending time points for each phoneme.

Next, we encoded *phonotactic probability* information in an appropriate two-dimensional vector (**Pt**)^33,39^. Probabilities were derived by means of the BLICK computational model^54^, which estimates the probability of a phoneme sequence to belong to the English language. This model is based on a combination of explicit theoretical rules from traditional phonology and a maxent grammar^104^, which find optimal weights for such constraints to best match the phonotactic intuition of native speakers. The phonotactic probability was derived for all phoneme sub-sequences within a word (ph_1..*k*_, 1 ≤ *k* ≤ *n*, where *n* is the word length) and used to modulate the magnitude of a phoneme onset vector (**Pt_1_**). A second vector was produced to encode the change in phonotactic probability due to the addition of a phoneme (ph_1..*k*_ - ph_1..*k-1*_, 2 ≤ *k* ≤ *n*) (**Pt_2_**).

Finally, a semantic dissimilarity descriptor was calculated for content words using word2vec^55,105^, a state-of-the-art algorithm consisting of a neural network for the prediction of a word given the surrounding context. In this specific application, a sliding-window of 11 words was used, where the central word was the output and the surrounding 10 words were the input. This approach is based on the “distributional hypothesis” that words with similar meaning occur in similar contexts, and it uses an artificial neural network approach to capture this phenomenon. This network has a 400-dimensional hidden layer that is fully connected to both input and output. For our purposes, the weights of this layer are the features used to describe each word in a 400-dimensional space capturing the co-occurrence of a content word with all others. In this space, words that share similar meanings will have a closer proximity. The semantic dissimilarity indices are calculated by subtracting from 1 the Pearson’s correlation between a word’s feature vector and the average feature vector across all previous words in that particular sentence (the first word in a sentence was instead correlated with the average feature vector for all words in the previous sentence). Thus, if a word is not likely to co-occur with the other words in the sentence, it should not correlate with the context, resulting in a higher semantic dissimilarity value. The semantic dissimilarity vector (**Sem**) marks the onset of content words with their semantic dissimilarity index.

### Computational model and data analysis

A single input event at time *t_0_* affects the neural signals for a certain time-window [*t_1_*, *t_1_*+*t*_win_], with *t_1_* ≥ 0 and *t_win_* > 0. Temporal response functions (TRF) were fit to describe the speech-EEG mapping within that latency-window for each EEG channel (TRF^106,107^). We did this by means of a regularized linear regression^58^ that estimates a filter that allows to optimally predict the neural response from the stimulus features (forward model; **Fig. 1C**). The input of the regression also included time-shifted versions of the stimulus features, so that the various time-lags in the latency-window of interest were all simultaneously considered. Therefore, the regression weights reflect the relative importance between time-latencies to the stimulus-EEG mapping and were here studied to infer the temporal dynamics of the speech responses (see **Figures 1** and **2**). Here, a time-lag window of 0–600 ms was used to fit the TRF models which is thought to contain most of the EEG responses to speech of interest. The reliability of the TRF models was assessed using a leave-one-out cross-validation procedure (across trials), which quantified the EEG prediction correlation (Pearson’s *r*) on unseen data while controlling for overfitting. Note that the correlation values are calculated with noisy EEG signal, therefore the *r*-scores can be highly significant even though they have low absolute values (*r* ~ 0.1 for sensor-space low-frequency EEG^32,64,100^).

Stimulus descriptors at the levels of acoustics, phonemes, phonotactics, and semantics were combined in a single TRF model fit procedure. This strategy was adopted with the goal of discerning EEG responses at different processing stages. In fact, larger weights are assigned to regressors that are most relevant for predicting the EEG. For example, a TRF derived with **Pt** alone could reflect EEG responses to phonotactics and phoneme onset. A TRF based on the combination of **Pt** and **Pon** would instead discern their respective EEG contributions, namely by assigning larger weights to **Pt** for latencies that are most relevant to phonotactics. Here, individual-subject TRFs were fit by combining **Env**, **Env’**, **Pvc**, **Pon, Pt**, and **Sem**(stimulus descriptor **ALL**). We also fit TRF models with an extended stimulus descriptor (**EXT**) including **Sgr**, **Env’**, **Phn**, **Pon, Pt**, and **Sem**, which provided us with a higher level of detail on spectrotemporal and phonological speech features, at the cost of higher dimensionality. The combined stimulus descriptor had 40 dimensions, which have to be multiplied by the number of time-lags (30 with sampling frequency 50Hz) to have the dimensionality of the TRF input. For this reason, we conducted all analysis on the reduced stimulus set **ALL**, while the **EXT** descriptor was used to assess spectrotemporal and phoneme TRFs.

The TRF weights constitute good features to study the spatio-temporal relationship between a stimulus feature and the neural signal. However, studying this relationship for multivariate speech descriptor, such as **Phn**, requires the identification of criteria to combine multiple dimensions of TRF weights. One solution to use the EEG prediction correlation values to quantify the goodness of fit for a multivariate TRF model. Here, we considered the relative enhancement in EEG prediction correlation when **Phn** was included in the model (using the **ALL** feature-set), thus allowing us to discern the relative contribution of phonetic features to the neural signal. This isolated index of phoneme-level processing was also shown to correlate with psychometric measures of phonological skills^35^. Additional analyses were conducted with a generic modelling approach^108^. Specifically, one generic TRF model was derived for each of the groups A, B, C, and L1 by averaging the regression weights from all subjects within the group. Then, EEG data from each left-out subject (whose data was not included in the generic models) was predicted with the four models. The four prediction correlations were used as indicators of how similar the EEG signal from a subject was to the one expected for each of the four groups, providing us with a simple classifier.

### Proficiency-level decoding

Support Vector Regression (SVR) with radial basis function kernel was used to decode the proficiency level of L2 participants. The output of the regression is the proficiency level, a continuous integer variable with six possible values corresponding with A1, A2, B1, B2, C1, and C2. The input of the SVR was the concatenation of 26 features derived from the TRF analysis described in the previous section. Specifically, we chose features based on the TRF weights (9 features), subject-specific EEG prediction correlations (5 features), and generic models EEG prediction correlations (12 features).

Each feature had multiple dimensions, such as EEG electrodes and time-latencies. A multilinear principal component analysis (MPCA) was performed to summarize each of them with a single value. The first component was retained for the TRFs corresponding to envelope, phoneme onsets, phonetic feature, phonotactics, and semantic dissimilarity. Based on previous TRF studies and our initial hypotheses, we complemented the result of this lossy compression by adding distinctive feature that summarize specific aspects of interest of the TRFs. For speech acoustics, we included information on the power spectrum of the TRF (the EEG responsiveness to 16 logarithmically-spaced sound frequencies) by collapsing the weights in TRF_ALL_ corresponding to **Sgr** across the time-latency dimension. MPCA was then run on the resulting values to quantify this spectral feature with a single value per subject. For phonotactics and semantic dissimilarity, the strength of the main TRF components was summarized by averaging the regression weights over selected time-windows and electrodes where they were strongest (80-140 and 300-700 for **Pt**, and 300-700 for **Sem**, at Fz, Cz, and Oz respectively).

EEG prediction correlations were calculated when TRF models were trained and tested on each participant separately (with leave-one-out cross-validation across recording blocks). This procedure provided us with a correlation score for each electrode, which was then summarized with a single value by running MPCA and retaining the first component. This procedure provided us with four features for EEG predictions based on **Env**, **Phn**, **Pt**, **Sem**. A fifth feature was derived by measuring the increase in EEG prediction correlations when **Phn** was included in the stimulus set by concatenating it with **Env’** and **Sgr**(**Phn_r_**). This subtraction was suggested to constitute an isolated measure of phoneme-level processing^32,35^. Finally, EEG signals from a subject were also predicted with TRF models fit on all other subjects, grouped in **A**, **B**, **C**, and **L1**, with the rationale that the EEG data from a given subject should be best predicted by TRF models from subjects of the same group. This approach, which has been referred to as average models or generic modelling approach^36,108^, provided each subject with a score for each group and for each feature of interest. Here we selected **Env’**, **Phn_r_**, and the concatenation of **Pt** and **Sem**. MPCA was then used for dimensionality reduction as for the other features, providing us with twelve features (4 groups and 3 predicting features).

SVR was used to decode the L2 proficiency level, for the binary classification L1 versus L2, or for the binary classification L1 versus C-level L2 with leave-one-out cross-validation. A backward elimination procedure was used to identify that optimal set of features that minimize the mean squared error (MSE) of the decoded proficiency levels. Specifically, starting from a set containing all the features, the regressor whose exclusion produced the larger decrease in MSE was removed at each step. This procedure continued as long as there was at least 5% improvement on the MSE score (please see **Supplementary Table 1** for a full list of features and information on the selected feature for the L2 decoding and L1 vs. L2 classification procedures).

### Statistical analysis

Statistical analyses were performed using two-tailed permutation tests for pair-wise comparisons. Correction for multiple comparisons was applied where necessary via the false discovery rate (FDR) approach. One-way ANOVA was used to assess when testing the significance of an effect over multiple (> 2) groups. The values reported use the convention *F*(*df*, *df_error_*). Greenhouse-Geisser corrections was applied when the assumption of sphericity was not met (as indicated by a significant Mauchly’s test). FDR-corrected Wilcoxon tests were used after the ANOVA for post-hoc comparisons.

## Acknowledgements

The authors would like to thank Michael Broderick for his help with the semantic dissimilarity analysis. The authors also thank Adam Soussana and Ghislain de Labbey for their help with a pilot version of this experiment.

## Author Contributions

The study was conceived by J.N., N.M., and G.D.L.; the experiments were designed by J.N. and B.K. programmed the tasks; J.N. collected the EEG data; G.D.L., N.M., J.N., J.Y. analyzed the data; G.D.L., J.N., N.M. wrote the first draft of the manuscript; J.Y., B.K., and S.S. edited the manuscript.

---

**Supplementary Figure 1.**
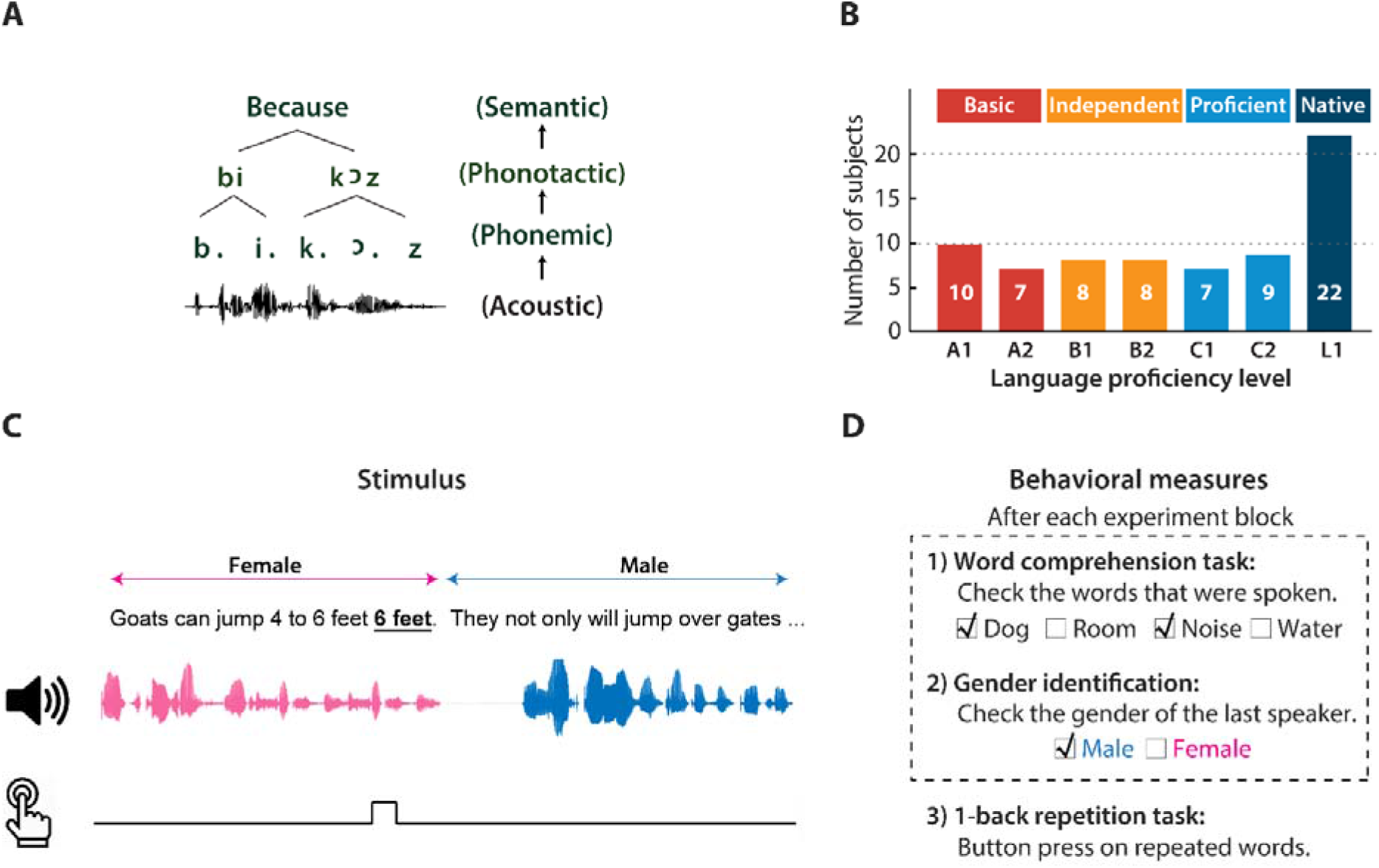
Experimental setup and behavioral results. (A) Selected part of the speech processing hierarchy, from low-level acoustics (bottom) to higher-level linguistic properties (top). (B) Demographic distribution of the proficiency levels. (C)Sentences spoken by a male and a female speaker were presented in alternation. Participants were asked to detect a one-back phrase repetition (2-4 words), which occurred 1-5 times per experimental block, by pressing a button during the experimental block. (D) After each block, participants were asked to identify words that were spoken during the block from a list of eight, and to indicate the gender of the speaker at the end of the block.

**Supplementary Figure 2.**
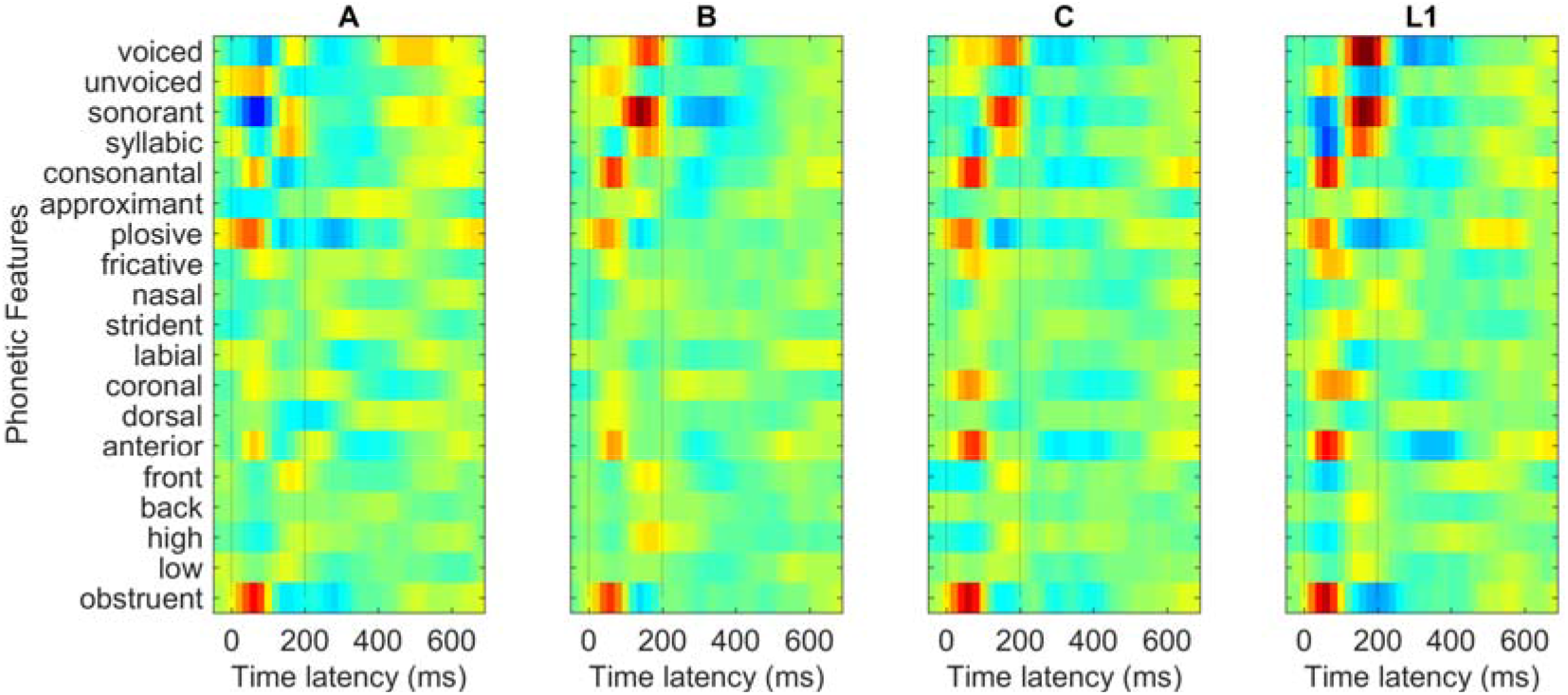
Average temporal response function weights across subjects at the electrode Cz and peri-stimulus time-latencies from 0 to 600 ms for each proficiency level. These TRF weights were used to produce the result in Figure 3A.

**Supplementary Table 1.**
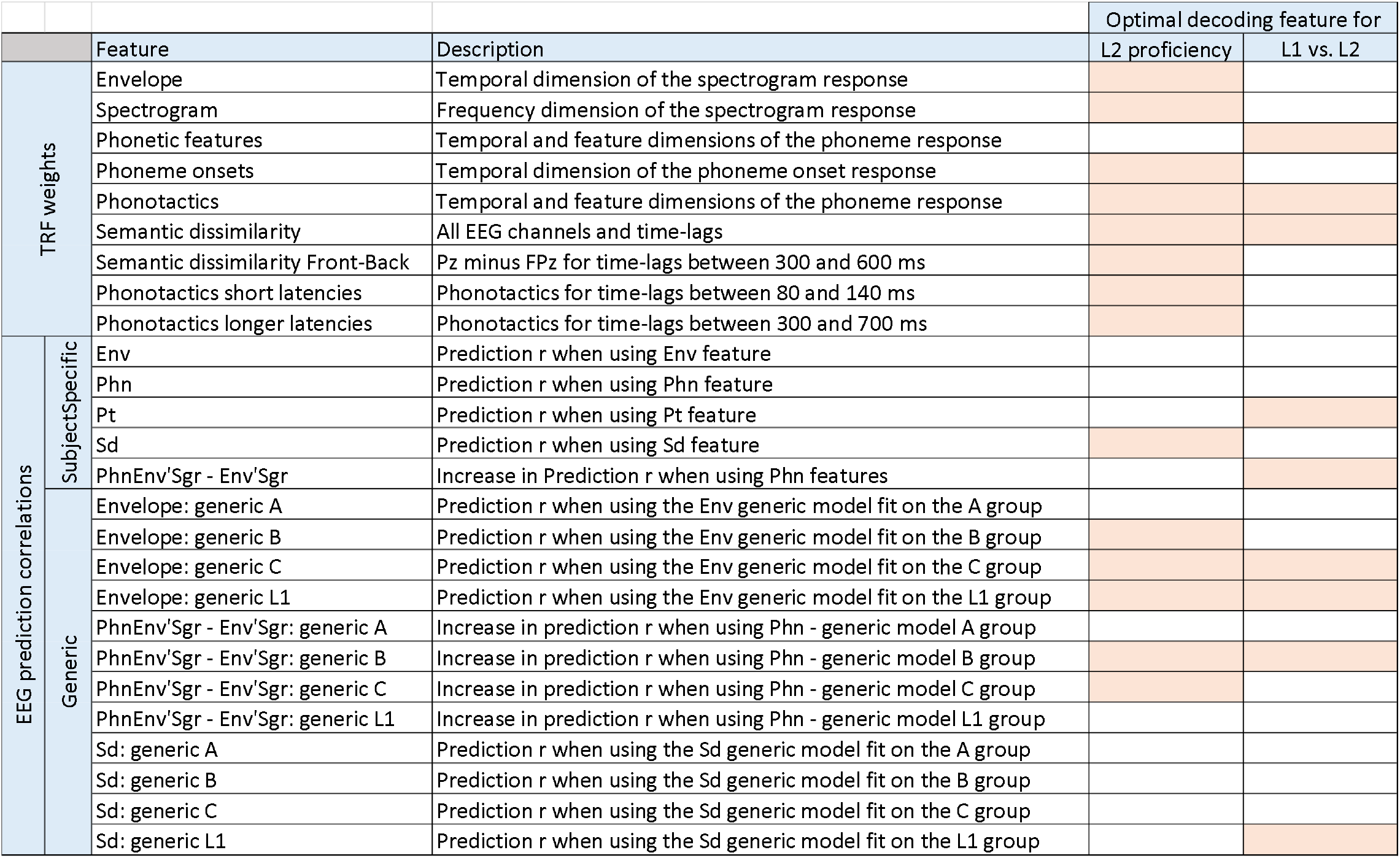
The 26 features used for the proficiency decoding and L1 vs. L2 classification analyses. Features are grouped into TRF weights, EEG prediction correlations when using subject-specific models, and EEG prediction correlations when using generic models that were averaged within a particular proficiency group. A backward elimination procedure was used for feature selection. Features that were selected for the decoding are indicated with a colored cell.

